# Modeling Uncertainty-Seeking Behavior Mediated by Cholinergic Influence on Dopamine

**DOI:** 10.1101/699595

**Authors:** Marwen Belkaid, Jeffrey L. Krichmar

**Affiliations:** Sorbonne Université, CNRS UMR 7222, Institut des Systèmes Intelligents et de Robotique, ISIR, F-75005 Paris, France; ETIS Laboratory, UMR 8051, Université Paris Seine, ENSEA, CNRS, Université de Cergy-Pontoise, F-95000 Cergy-Pontoise, France; Department of Cognitive Sciences, University of California, Irvine, Irvine, CA 92697, USA; Department of ComputerScience, University of California, Irvine, Irvine, CA 92697, USA

## Abstract

Recent findings suggest that acetylcholine mediates uncertainty-seeking behaviors through its projection to dopamine neurons – another neuromodulatory system known for its major implication in reinforcement learning and decision-making. In this paper, we propose a leaky-integrate-and-fire model of this mechanism. It implements a softmax-like selection with an uncertainty bonus by a cholinergic drive to dopaminergic neurons, which in turn influence synaptic currents of downstream neurons. The model is able to reproduce experimental data in two decision-making tasks. It also predicts that i) in the absence of cholinergic input, dopaminergic activity would not correlate with uncertainty, and that ii) the adaptive advantage brought by the implemented uncertainty-seeking mechanism is most useful when sources of reward are not highly uncertain. Moreover, this modeling work allows us to propose novel experiments which might shed new light on the role of acetylcholine in both random and directed exploration. Overall, this study thus contributes to a more comprehensive understanding of the roles of the cholinergic system and its involvement in decision-making in particular.

## 1 Introduction

Animals constantly face uncertainty due to noisy and incomplete information about the environment. From the information-processing perspective, uncertainty is typically considered a burden, an issue that has to be resolved for the animal to behave correctly [Cohen et al., 2007; Rao, 2010]. In the framework of reinforcement learning, for example, to allow optimal exploitation and outcome maximization, agents must explore the environment and gather information about action–outcome contingencies [Sutton and Barto, 1998; Rao, 2010].

The neural mechanisms driving the decision to perform actions with uncertain outcomes are still poorly understood. In contrast, the processes by which individuals learn to perform successful actions have been extensively studied. Notably, the dopaminergic system is thought to play a key role in these processes, both in the learning- and in the motivation-related aspects [Schultz, 2002; Berridge, 2012; Berke, 2018]. Moreover, studies have reported dopaminergic activities that are correlated with the uncertainty of reward [Fiorillo et al., 2003; Linnet et al., 2012].

Another neuromodulatory system which has been largely implicated in the processing of novelty and uncertainty is the cholonergic system. For instance, Yu and Dayan [2005] suggested that acetylcholine (ACh) suppresses top-down, expectation-driven information relative to bottom-up, sensory-induced signals in situations of expected uncertainty, i.e. when expectations are known to be unreliable. Additionally, Hasselmo [1999, 2006] proposed that the level of ACh in the hippocampus determines whether it is encoding new information or consolidating old memories. The cholinergic system also interacts with the dopaminergic system. In particular, there are cholinergic projections onto neurons in the ventral tegmental area (VTA), one of the two major sources of dopamine (DA) in the brain [Avery and Krichmar, 2017; Scatton et al., 1980]. In a recent study, Naudé et al. [2016] provided evidence that these projections might mediate the motivation to select uncertain actions.

The softmax rule, where the probability of choosing an action is a function of its estimated value, is generally thought to be a good model of human [Daw et al., 2006] and animal [Cinotti et al., 2019] decision-making. But Naudé et al. [2016] showed that the decisions made by wild-type (WT) mice exhibited an uncertainty-seeking bias and followed a softmax function which included an uncertainty bonus. In contrast, mice lacking the nicotinic acetylcholine receptors on the dopaminergic neurons in VTA showed less uncertainty-seeking behaviors and their decisions rather followed the standard softmax rule.

In neural networks, decision-making processes are generally modeled using competition mechanisms [Rumelhart and Zipser, 1985; Carpenter and Grossberg, 1988]. Such mechanisms can constitute a neural implementation of the softmax rule. In particular, Krichmar [2008] proposed a model where neurotransmitters act upon different synaptic currents to modulate the network’s sensitivity to differences in input values, much like the temperature parameter in the softmax model [Sutton and Barto, 1998]. In this paper, we propose a new version of this model using leaky-integrate-and-fire neurons and integrating an uncertainty bonus. We use this model, in comparison with three alternative models, to verify a set of hypotheses about how cholinergic projections to dopaminergic VTA neurons in mediate uncertainty-seeking. We then perform additional experiments to assess the interest of such a mechanism for animals foraging in volatile environments. These simulations suggest that ACh effects behavior by translating uncertainty into a source of motivation thus driving exploratory behaviors.

## 2 Background

### 2.1 Dopamine

Dopamine (DA) is involved in decision-making through its role in reward processing and motivation [Schultz, 2002; Berridge, 2012]. The largest group of dopaminergic neurons is found in the ventral tegmental area (VTA) [Scatton et al., 1980]. It projects to the basal ganglia (BG), in particular to the striatum, but also to the frontal cortex. The substantia nigra is also an important source of dopamine in the BG.

There is strong evidence of the role of dopamine in the learning of the value of actions, stimuli and states of the environment. In this context, Schultz and colleagues hypothesized that the activity of DA neurons encodes a reward prediction error [Schultz, 2002]. Indeed, phasic dopaminergic activities show strong correlations with an error in the prediction of conditioned stimuli after Pavlovian learning. Moreover, Berridge and colleagues suggest that DA is essential for “incentive salience” and “wanting”, i.e. for motivation [Berridge and Kringelbach, 2008; Berridge, 2012]. For instance, DA deprived rats are unable to generate the motivation arousal necessary for ingestive behavior and can starve to death although they are able to move and eat [Ungerstedt, 1971]. However, dopamine has also been suggested to signify novelty, which may be related to an uncertainty signal [Redgrave and Gurney, 2006]. In summary, the dopaminergic system seems to implement a series of mechanisms that reinforce and favor stimuli and actions that have been rewarding in the past, or that may be of interest in the future.

### 2.2 Acetylcholine

Acetylcholine (ACh) originates from various structures in the brain: the laterodorsal tegmental (LDT) and the pedunculopontine tegmental (PPT) mesopontine nuclei projecting to the VTA and other nuclei in the brainstem, basal forebrain and basal ganglia [Mena-Segovia, 2016]; the medial septal nucleus mainly targeting the hippocampus; and the nucleus basalis in the basal forebrain mainly acting on the neocortex [Baxter and Chiba, 1999]. In addition, striatal interneurons provide an internal source of ACh in the BG.

ACh has been largely implicated in the processing of novelty and uncertainty. Significant research highlighted this role in the septohippocampal cholinergic system for instance. In this case, novelty detection increases the level of septal ACh: novel patterns elicit little recall which reduces hippocampal inhibition of the septum and allows ACh neurons to discharge [Meeter et al., 2004]. In addition, Hasselmo [1999, 2006] proposed that high and low levels of ACh in the hippocampus – during active waking on the one hand, and quiet waking and slow-wave sleep on the other hand – respectively allow encoding new information and facilitate memory consolidation. Similarly, higher activity of the cholinergic neurons in the tegmentum and nucleus basalis have been shown to be associated with cortical activation during waking and paradoxical sleep [Jones, 2005] – a sleep phase physiologically similar to waking states. Thus, various computational models of the cholinergic system have focused on its role in learning and memory [Hasselmo, 2006; Pitti and Kuniyoshi, 2011; Grossberg, 2017].

A complementary theory was developed by Yu and Dayan [2005] suggesting that acetylcholine suppresses top-down, expectation-driven information relative to bottom-up, sensory-induced signals in situations of expected uncertainty, i.e. when expectations are known to be unreliable. To illustrate their theory, the authors modeled the so-called Posner task. Posner [1980] proposed this paradigm to study attentional processes. Typically, a cue is presented to the participants, followed by a target stimulus. Posner [1980] showed that individuals respond more rapidly and accurately on correctly cued trials (i.e. cue on the same side as the target) than on incorrectly cued trials (i.e. cue on opposite side). The difference in response time between valid and invalid trials is termed *validity effect* (VE). The model proposed by Yu and Dayan [2005] reproduced the results obtained by Phillips et al. [2000] which showed in rats experiments that VE varies inversely with the level of ACh which was manipulated pharmacologically. Additionally, ACh has been hypothesized to set the threshold for noadrenergic signaling of unexpected uncertainty [Yu and Dayan, 2005] which calls for more exploration by counterbalancing DA-driven exploitation [Cohen et al., 2007].

## 3 Methods

### 3.1 Bandit task

The experiment reported by Naudé et al. [2016] was a 3-armed bandit task adapted for mice. The setup was an open-field in which three target locations were associated with a certain probability of rewards (**Figure 1A**), which was delivered through intracranial self-stimulation (ICSS). Mice could not receive two consecutive ICSS at the same location. Thus, each time they were at a target location, they had to choose the next target among the two remaining alternatives. As in a classical bandit task, this is referred to as a gamble. Since the outcome is binary (i.e. reward delivered or not), the expected uncertainty was represented by the variance *p*(1 − *p*) of Bernoulli distributions (**Figure 1B**).

**Figure 1:**
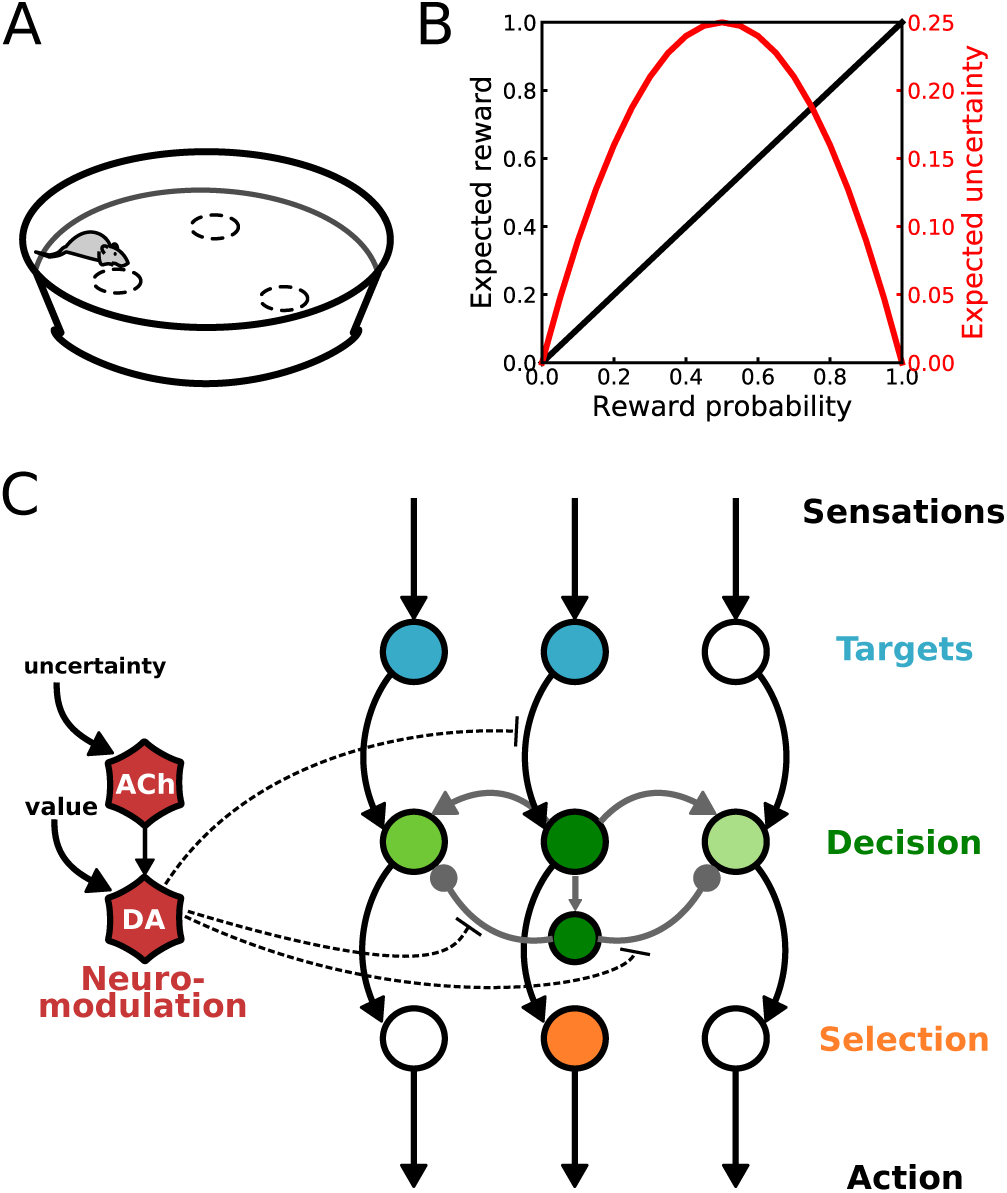
Bandit task and core model. **A)** Task setup used by Naudé et al. [2016]. **B)** Expected reward and uncertainty as a function of reward probability in this task. **C)** Neural network model of decision-making. (see text for description).

Naudé et al. [2016] used this task to study the influence of uncertainty on decision-making, and more specifically on the dopaminergic activity under the influence of cholinergic projections. Notably, they showed that while wild type (WT) mice exhibited uncertainty-seeking behavior in their task, such behaviors were suppressed in mice with deleted nicotinic acetylcholine receptors in the dopaminergic neurons in VTA (hereafter KO mice).

### 3.2 Neural Network Model of Uncertainty Seeking

We model the decision-making process involved in this task using an artificial neural network (**Figure 1C**). This network has three channels, each corresponding to one of the targets. Similar to Krichmar [2008], the competition takes place in a decision layer were neurons have lateral excitatory and inhibitory connections in addition to extrinsic input from downstream layers. Neuromodulatory signals driven by the cholinergic and dopaminergic representative neurons modulate this competition. When the dopaminergic activity is low, the low signal-to-noise ratio in decision neurons leaves room for exploration. However, strong dopaminergic activity amplifies the efficacy of extrinsic input connections and those of inhibitory interconnections in order to achieve exploitative decisions. This neuromodulation thus implements a neuronal equivalent to the softmax decision policy based on the value of the target. Moreover, the cholinergic activity increases the firing of DA and introduces an uncertainty bias in the softmax-like neuromodulation of the competition. In the bandit task simulations, the value and uncertainty signals are manually provided to the model (see Section 3.2.1).

All neurons are leaky-integrate-and-fire (LIF) neurons. The change in the membrane potential *V* is represented as follows:

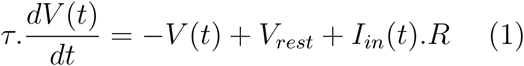

where *τ* is the time constant, *V*_*rest*_ is the resting potential, *R* the resistance of the membrane, and *I*_*in*_ the input current:

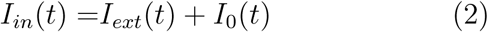

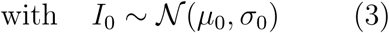

where *I*_*ext*_ and *I*_0_ are respectively extrinsic and background input currents. The latter is modeled as a Gaussian distribution 𝒩(*µ*_0_, *σ*_0_) and accounts for spontaneous activities as well as possible other extrinsic inputs which are not specifically modeled here. When the membrane potential is higher than a threshold *V*_*th*_, the neuron fires, i.e. the potential rises to *V*_*spike*_ then decreases to *V*_*rest*_ and a current *I*_*out*_ is transmitted to post-synaptic neurons:

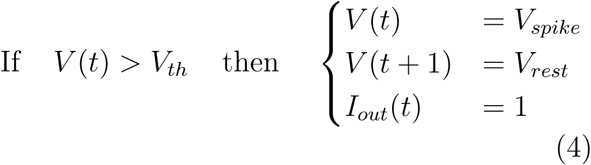

A single trial of the experiment consists of a decision made between two target locations. For simplicity, neurons of the target identification layer (in blue in **Figure 1A**) were tuned such that they fire every two iterations (a spike is followed by a refractory period) with random initialization. Only the two target neurons corresponding to the current options are activated in each trial.

Similarly to the model used by Krichmar [2008], the extrinsic input current of the decision neurons 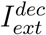 (in green in **Figure 1A**) is defined as follows:

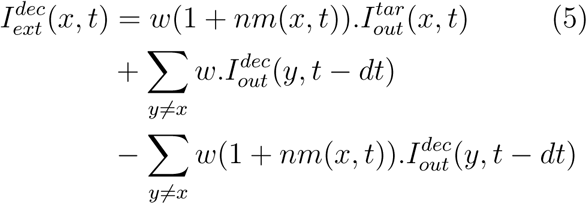

where *x* ∈{*A, B, C*} corresponds to the gambling options, *w* is a synaptic weight and *nm* is a neuromodulation factor that we will define below.

As for selection neurons (in orange in **Figure 1A**), the extrinsic input current is simply 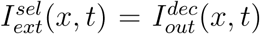. This layer implements a winner-takes-all readout of the decision. The first spike corresponds to the network’s decision.

In this model, decisions are modulated by the dopaminergic system. To account for the difference between wild type (WT) mice and mice in which nicotinic achetylcholine receptors in dopamine neurons were removed (KO) reported by Naudé et al. [2016], we defined two variants of the neuromodulation component: the WT and KO models.

#### 3.2.1 WT model

The ability to learn the reward probability and maximize the outcome is thought to be mediated by the dopaminergic system. Thus, in our model, DA activity is a function of the targets value *v*(*x*) representing the reward probability. Besides, Naudé et al. [2016] observed an uncertainty-driven motivation in WT mice in addition to the motivation to maximize reward by choosing the target with highest reward probability. They showed that this uncertainty-seeking behavior was dependent upon the cholinergic projections to DA neurons, which also modulate the dopaminergic activity. Since the expected uncertainty is thought to be encoded by ACh neurons, in our model, ACh activity is determined by the reward uncertainty *u*(*x*) which we represent as the variance of a Bernouilli distribution *v*(*x*)(1 *- v*(*x*)) (Figure 1B). We define *I*_*u*_ = ∑_*x∈*(*O*)_ *u*(*x*) as an input current generated by the overall expected uncertainty in the current trial, and *I*_*v*_ = ∑_*x*∈(*O*)_ *v*(*x*) an input current generated by the overall expected rewards in the current trial. Thus, the activity in neuromodulation network is determined by the following equations:

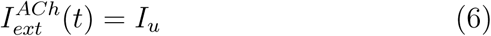

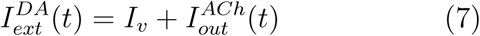

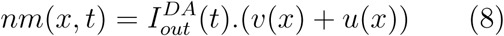

Introducing the output current of the ACh neuron as an input to the DA neuron is consistent with the increase of dopaminergic activity observed in the presence cholinergic receptors [Graupner et al., 2013; Naudé et al., 2016]. In the bandit task, *v*(*x*) and *u*(*x*) of each target were manually fixed for simplicity. But in the foraging task, these variables were estimated by the model (see Section 3.5).

#### 3.2.2 KO model

Naudé et al. [2016] showed that uncertainty-seeking was removed in KO mice. Their decisions were rather exploitative, similarly to a classical softmax policy. Thus, in this variant of the model, the cholinergic effect on decision is eliminated and the dopaminergic activity only depends on reward probabilities:

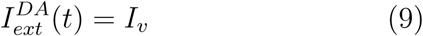

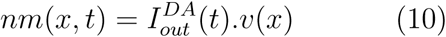

### 3.3 Alternative models

The proposed model assumes that: i) uncertainty is processed by ACh, and that ii) cholinergic projections to DA neurons not only increase the latter’s firing rate but also the neuromodulation of the action selection process. To further validate these hypotheses, we tested three alternative models – all including a WT and a KO variants – introducing the following changes.

#### Alternative model 1

In this model, ACh simply increases DA firing rate independently from uncertainty. It is set to fire at a similar rate as previously using a constant input. But, uncertainty is neither processed by ACh nor DA. Hence, there is no difference in the neuromodulation term between the WT and KO variants, both using the form in Equation (10). The only difference between the WT and KO variants is whether ACh activates DA.

#### Alternative model 2

In this model, the ACh neuron also has a constant input independent from reward uncertainty. However, uncertainty is processed by DA neurons. Hence, there is no difference in the neuromodulation term between the WT and KO variants, this time both using the form in Equation (8). The only difference between the WT and KO variants is again whether ACh activates DA:

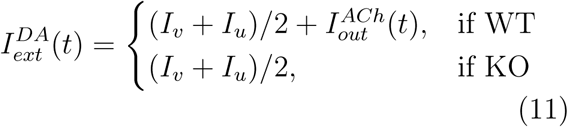

#### Alternative model 3

In this model, everything is similar to the WT model with one exception: ACh projections only increase DA firing rate. Hence, there is again no difference in the neuromodulation term between the WT and KO variants, this time both using the form in Equation (10). The only difference between the WT and KO variants is again whether ACh activates DA. This model differs from Alternative model 1 in that ACh firing is not driven by a constant input but rather by the estimated uncertainty of reward.

### 3.4 Foraging task

Naudé et al. [2016] did an additional experiment with a dynamic setup simulating a volatile environment. More specifically, in each session, two of the targets were rewarding 100% of time while the remaining one was not. The non-rewarding target changed from one session to another (**Figure 4A**), which required that animals detect the change of rule and learn the new reward probabilities.

**Figure 2:**
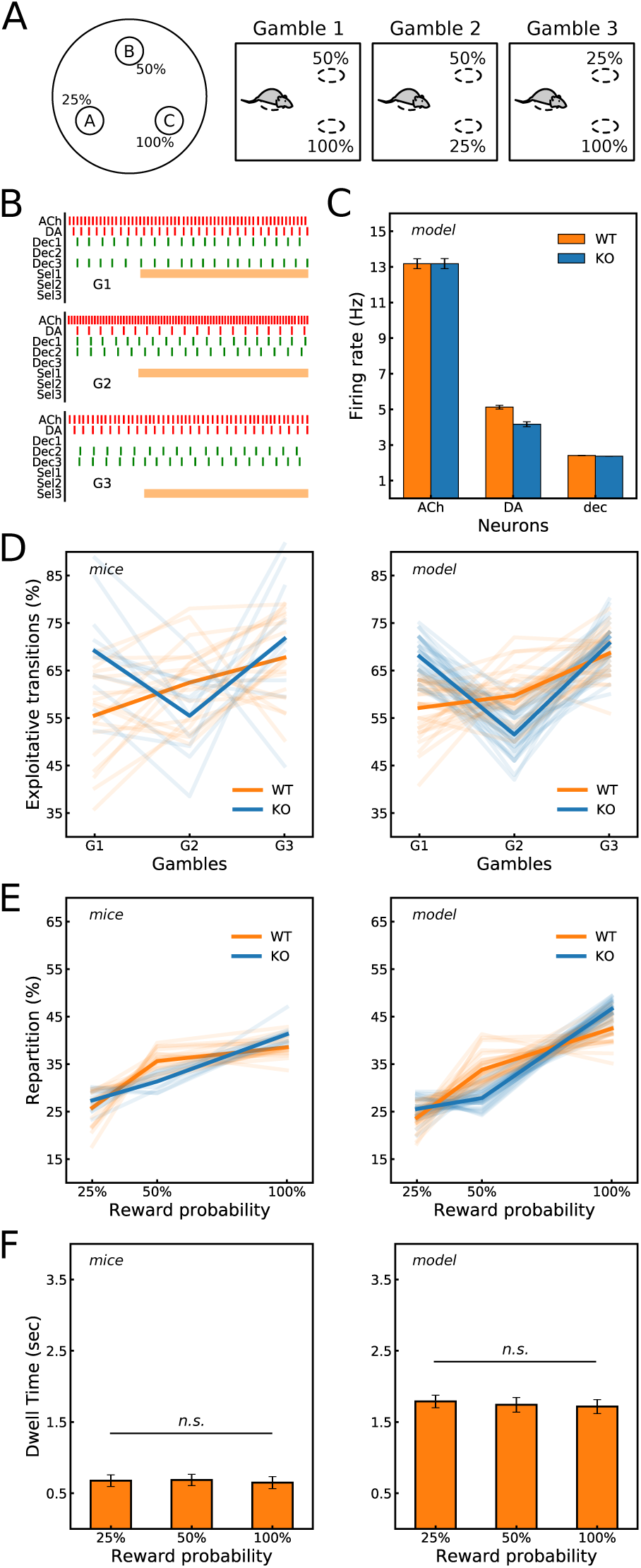
The proposed model reproduces mice behavior. **A)** Schematic illustration of the task setup and the three possible gambles. **B)** Example of spike trains generated by the model. **C)** Mean firing rate produced by the WT and KO versions of the model. **D)** Percentage of exploitative transitions (i.e. choosing the option with the highest reward probability) in each gamble. WT and KO mice (*Left*) had distinct profiles, which the WT and KO models (*Right*) were able to reproduce. **E)** Percentage of targets selection as a function of their reward probability. The model (*Right*) also reproduced the repartition of choices exhibited by mice (*Left*). **F)** Dwell time (i.e. time to decision) was also similar between targets with our model (*Right*), like in mice (*Left*). Mice results were plotted with data from Naudé et al. [2016].

**Figure 3:**
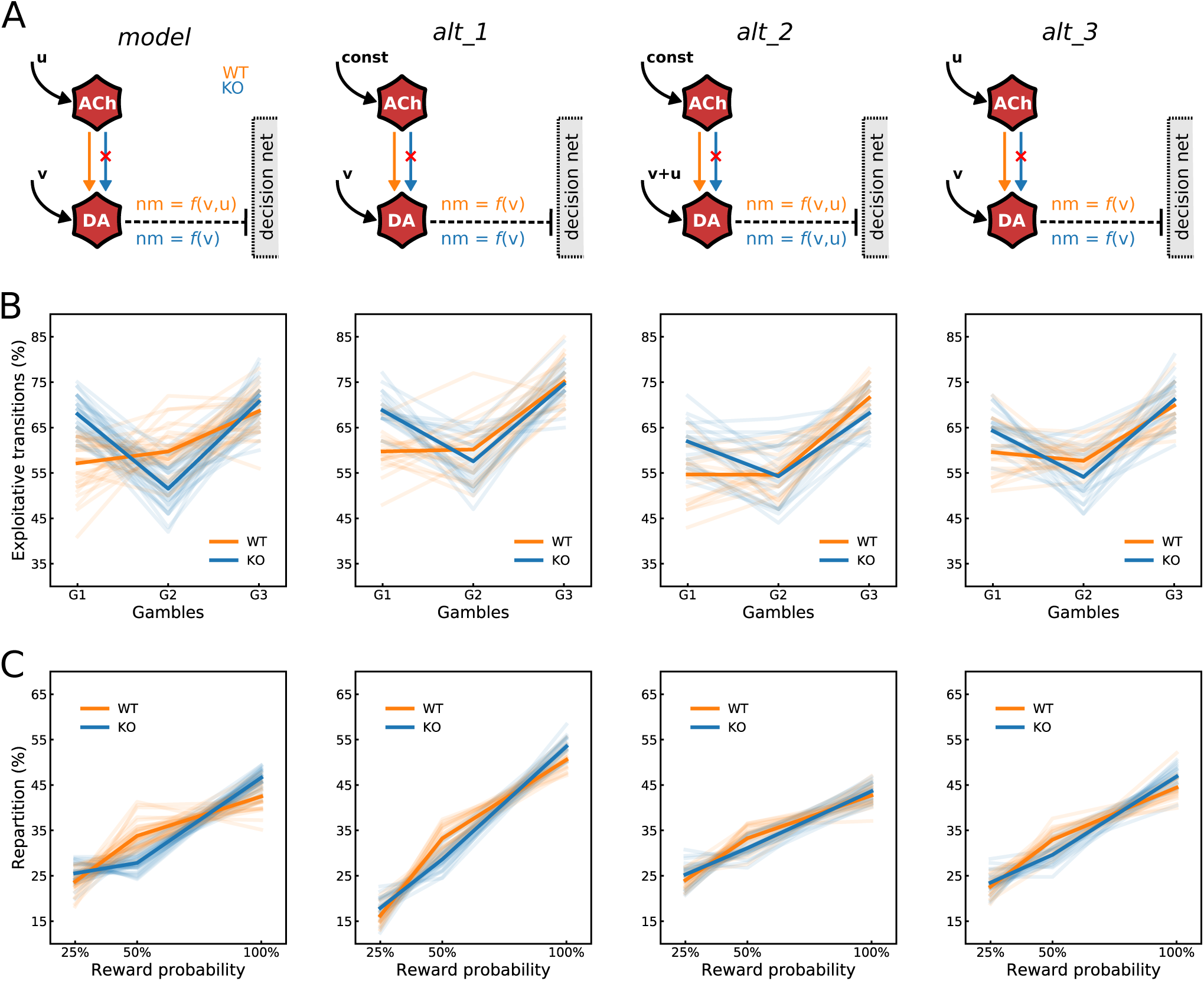
Alternative models fail to fully reproduce mice behavior. **A)** Schematic illustration of the differences in the neuromodulatory component between the core model and the three alternative models. **B)** Percentage of exploitative transitions. **C)** Percentage of targets selection as a function of their reward probability across gambles. In **B)** and **C)**, the alternative models did not fit the profiles observed in WT and KO mice.

**Figure 4:**
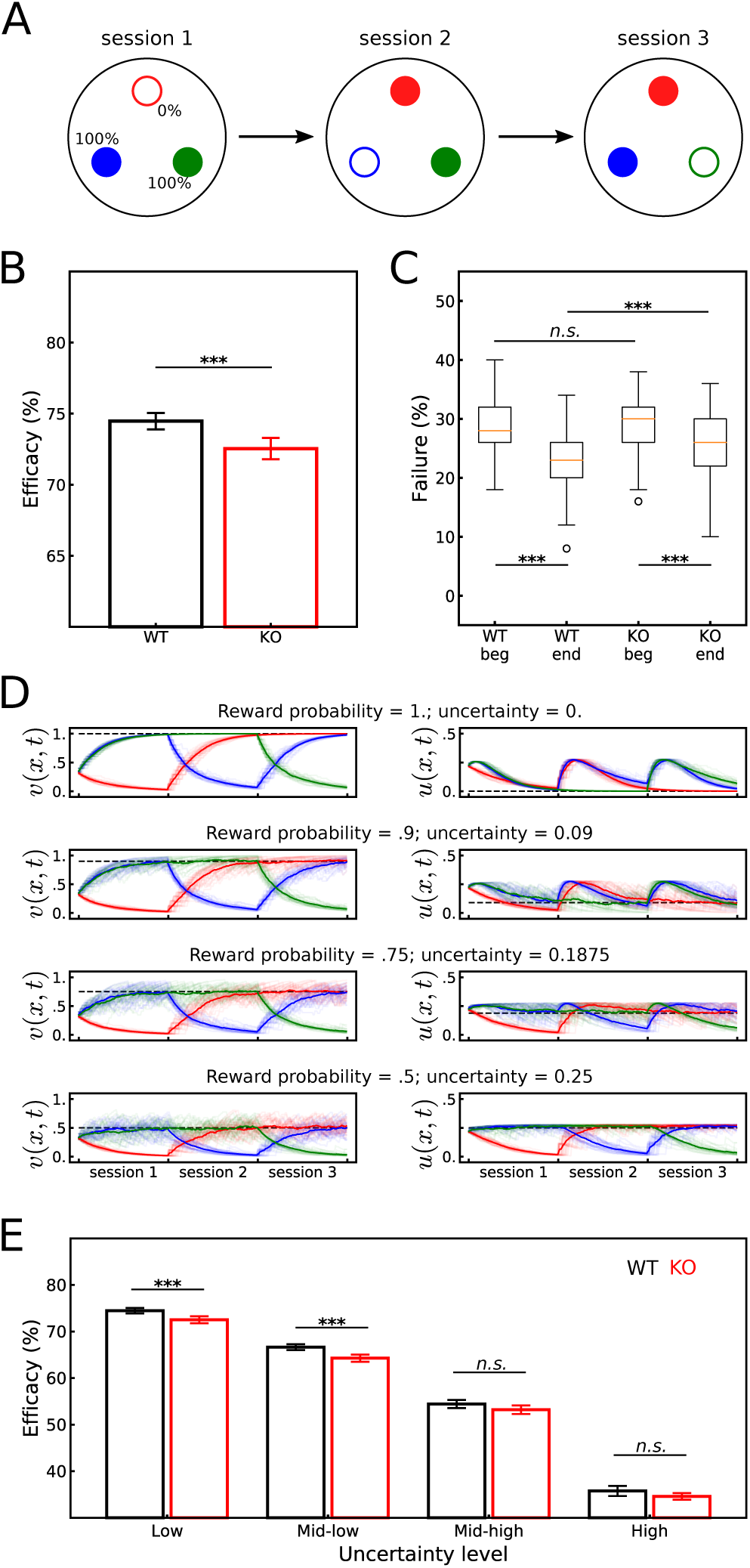
Model’s results and predictions in the foraging task. **A)** Schematic illustration of the dynamic setup consisting of three sessions. Full circles indicate the two rewarding targets and empty circles indicate the non-rewarding target. **B)** Higher foraging efficacy with the WT model than KO model. Efficacy is defined as the success rate, i.e. the average proportion of rewarded choices. **C)** Failure rate in the beginning and in the end of sessions shows a decrease for both WT and KO models but is lower for WT. **D)** Reward prob-ability *v* and uncertainty *u* were correctly estimated by the model throughout sessions. Dashed lines indicate the correct values. **E)** The model predicts that the difference in foraging efficacy between WT and KO mice vanishes in situations where the reward uncertainty is high. *** p < 0.001, n.s. not significant at p > 0.05.

### 3.5 Learning task statistics

In Equations (6 - 10), the expected reward probability *v* and uncertainty *u* for each target were manually fixed. But to model the dynamic foraging task, these statistics about the environment outcomes could no longer be hardwired and had to be learned by trial-and-error.

To learn the expected reward probabilities, we used the Rescorla-Wagner rule [Rescorla and Wagner, 1972]:

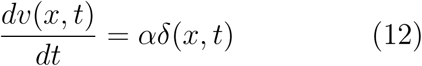

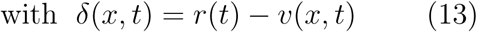

where *r* is the reward function, equal to 1 when a reward is obtained, and to 0 otherwise.

Additionally, the reward uncertainty could be estimated as follows [Balasubramani et al., 2014; Naudé et al., 2016]:

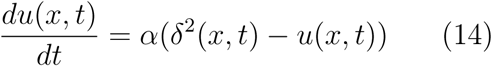

The hyperparameter *α* was set to 0.1.

### 3.6 Model fitting

The hyperparameters *τ, V*_*spike*_, *V*_*th*_, *V*_*rest*_, *µ*_0_ and *σ*_0_ are common to all neurons and were set manually so as to determine the dynamics of the network (see values in **Table 1**). The values of *R*^*ach*^ and *R*^*da*^, i.e. the membrane resistance in ACh and DA neurons respectively, was accordingly set to match the mean firing rate reported by Naudé et al. [2018]. We did not have data on the firing rate of ACh neurons in the mesopontine nuclei but the obtained firing rate is within the range that has been reported for other areas.

**Table 1:** Hyperparameter values (when set) and ranges (when optimized). The hyperparameters *τ, V*_*spike*_, *V*_*th*_, *V*_*rest*_, *µ*_0_ and *σ*_0_ were chosen manually. *R*^*ach*^ and *R*^*da*^ were set to fit experimentally observed firing rates of DA neurons. *R*^*dec*^, *R*^*sel*^ and *w* were optimized through a grid search.

| Hyperparameter | Value |
| --- | --- |
| $\tau$ | 20 |
| $V_{spike}$ | 5 |
| $V_{th}$ | 1 |
| $V_{rest}$ | -2 |
| $\mu_0$ | 0.15 |
| $\sigma_0$ | 0.05 |
| $R^{ach}$ | 60 |
| $R^{da}$ | 5.5 |

**Table 1: Hyperparameter values (when set) and ranges (when optimized).**
| Hyperparameter | Range | Step |
| --- | --- | --- |
| $R^{dec}$ | [10, 60] | 1 |
| $R^{sel}$ | [5, 16] | 1 |
| $w$ | [0, 1] | 0.05 |

The values of *R*^*dec*^, *R*^*sel*^ and *w*, i.e. respectively the membrane resistance in the decision and the selection layers and the baseline synaptic weight in the lateral connection within the decision layer, were optimized using a grid search (see ranges listed in **Table 1**) to fit the proportion of exploitative choices observed by Naudé et al. [2016] in WT and KO mice. The models’ results were averaged over 30 runs comprising 300 trials each and the fitness score *S* was calculated as follows:

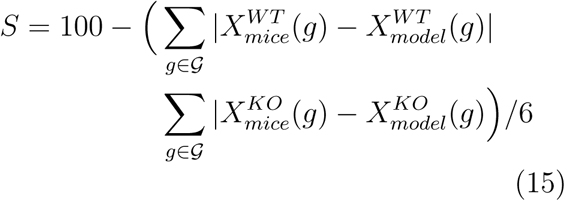

where *X* is the average proportion of exploitative choices and *𝒢* is the set of gambles.

## 4 Results

### 4.1 Bandit task

In this task reported by Naudé et al. [2016], animals had to make binary choices (called *gambles)* among the remaining two out of three target locations that were set to deliver rewards with probabilities *P* = 25%, 50% and 100% respectively (**Figure 2A**). We modeled this decision-making process with a neural network (**Figure 1C**). The hyperparameters determining the dynamics of the model were first manually set to match the mean firing rate reported by Naudé et al. [2018] in DA neurons *in vivo* (**Figure 2B** and **C**). The remaining hyperparameters of the model were optimized to fit the proportion of exploitative choices observed by Naudé et al. [2016] in WT and KO mice (**Figure 2D**). As a result, the model reproduced experimental data (**Figure 2D**). Notably, the two groups had distinct profiles which respectively correspond to uncertainty bonus and standard softmax decision rules.

Interestingly, the WT and KO models also reproduced the repartition of choices among targets (i.e. overall percentage of times each target was selected across trials) that was observed by Naudé et al. [2016] (**Figure 2E**).

As with the softmax rule, KO mice and the corresponding model selected targets proportionally to their probability of reward whereas WT mice and the corresponding model exhibited a bias in favor of uncertainty in the case of reward probability 50%. Additionally, like in mice, there was no difference between the targets in terms of dwell time (i.e. time to make a decision, calculated in the model as the time of the first spike in the trial). In other words, there was no effect of the reward probability on the decision time (*H* = 4.70, *p* = 0.09, Kruskal-Wallis test; **Figure 2F**). Importantly, these two criteria (repartition of choices and dwell time) were not explicitly optimized by the model fitting procedure.

To further validate our model, we tested three alternative models introducing two types of changes in the neuromodulatory component: i) uncertainty could be either encoded by dopamine directly or not taken into account at all, ii) the absence of ACh receptors on DA only affected the latter’s firing but not the neuromodulatory effect (**Figure 3A**; see ‘Methods’ for more detailed explanation). Upon optimization, none of the alternative models was able to fully fit the behavioral data. Indeed, the fitness scores (calculated using Equation (15)) for these alternative models were significantly lower than our model’s (model versus alt1, *t*(29) = 3.38, *p* = 0.0013, model versus alt2, *t*(29) = 5.63, *p* = 10^*-*6^, model versus alt3, *t*(29) = 3.35, *p* = 0.0014, *t*-test; **Table 2**). Fitness scores quantify the ability of the WT and KO variants of a model to fit the proportion of exploitative transitions made by the corresponding group of mice. Lower scores can be explained by the fact that WT variants of the alternative models did not follow the same linear increase from gamble 1 to 3 (**Figure 3B**). Moreover, qualitatively, the differences in exploitative transitions and probability of selection of each target between the WT and KO model were smaller than with our model **(Figure 3B and C)**.

**Table 2:** Model fitting results. Optimized parameters and fitness scores. The proposed model has the highest score. *R*^*dec*^ and *R*^*sel*^ are the membrane resistance in the decision layer and the selection layer respectively. Stars indicate the results of a *t*-test comparison between the model’s score to each of the alternative models’ score: ** p < 0.01, *** p < 0.001.

| | $R^{dec}$ | $R^{sel}$ | $w$ | Score |
| --- | --- | --- | --- | --- |
| *model* | 12 | 12 | 0.7 | **96.19** |
| *alt 1* | 59 | 5 | 1 | 94.95** |
| *alt 2* | 43 | 7 | 0.6 | 94.72*** |
| *alt 3* | 10 | 13 | 0.8 | 95.06** |

### 4.2 Foraging task

We also tested our model in a foraging task where only two of the targets were rewarding. The non-rewarding target changed from one session to another (**Figure 4A**). In such a volatile environment, animals must detect the changes in reward probabilities and adapt their decisions accordingly. We initially tested a setup in which rewarding targets had 100% probability in the original experiments [Naudé et al., 2016]. In line with the experimental results, we found that the KO model had a lower foraging efficacy (i.e. global reward rate) than the WT model (WT versus KO: *t*(29) = −3.92, *p* = 0.0002, *t*-test; (**Figure 4B**). We split the sessions in half to analyze the model’s behavior more closely (**Figure 4C**). The WT and KO models had similar failure rates in the beginning of sessions (WT versus KO, *U* = 4249.0, *p* = 0.566, Mann-Whitney test), and both significantly reduced their failure rates at the end of session (beginning versus end of session for WT, *T* = 854.5, *p* = 6.10^*-*11^, for KO, *T* = 854.5, *p* = 5.10^*-*5^, Wilcoxon test). However, the rate of failure was significantly lower at the end of session for the WT model (WT versus KO, *U* = 5695.5, *p* = 2.10^*-*6^, Mann-Whitney test), suggesting that the KO model adapted more slowly to condition changes.

To assess how robust was this effect on foraging efficacy, we further tested similar setups where reward probability in the two rewarding targets were lower (but still equal) resulting in higher uncertainty: p=90%, 75% and 50% probability of reward corresponding to mid-low, mid-high and high uncertainty. The model successfully estimated the expected reward probability *v* and uncertainty *u* (**Figure 4D**; see Equations (12 - 14)). While the foraging efficacy was still higher for the WT model with reward probability 90% (WT versus KO: *t*(29) = −4.64, *p* = 2.10^*-*5^, *t*-test; **Figure 4E**), the difference was no longer significant with a probability of 75% (WT versus KO: *t*(29) = −1.89, *p* = 0.06, *t*-test; **Figure 4E**) and 50% (WT versus KO: *t*(29) = −1.73, *p* = 0.08, *t*-test; **Figure 4E**). Overall, these results demonstrate the interest of the uncertainty-seeking behaviors mediated by the cholinergic projections to VTA dopaminergic neurons but suggest that the scope of such an adaptively advantageous mechanism is limited to situations where the uncertainty is low.

## 5 Discussion

Prominent theories about the role of acetylcholine hold that it helps control the balance between the storage and update of memory [Hasselmo, 1999] and between top-down expectation-driven and bottom-up stimulus-driven attention [Yu and Dayan, 2005; Cohen et al., 2007; Avery et al., 2012]. Accordingly, most computational models of this neuromodulator at the functional level focus on memory- and attention-related functions [Hasselmo, 2006; Pitti and Kuniyoshi, 2011; Grossberg, 2017; Yu and Dayan, 2005; Avery et al., 2012]. In this paper, we targeted another aspect of the cholinergic action which was highlighted in recent experimental studies [Naudé et al., 2016, 2018]. These studies suggest that, through its projections to dopaminergic neurons in the ventral tegmental area, acetylcholine also promotes exploratory uncertainty-seeking behaviors. In other words, that the neuromodulator participates in the process by which individuals decide to perform actions associated with uncertain outcomes.

We modeled this process using a decision-making neural network under the influence of cholinergic and dopaminergic modulation. This model is based on the following hypotheses: a) dopamine encodes the estimated value, b) dopamine modulates the decision-making network such as to implement a softmax-like rule, c) acetylcholine encodes the estimated uncertainty, d) acetylcholine increases dopamine firing, e) acetylcholine introduces an uncertainty bonus in the softmax-like decision rule. The model was tested in two decision-making tasks – bandit task and foraging task – and successfully reproduced the behavioral results reported by Naudé et al. [2016]. In addition, the model fitted t he e xperimental d ata f rom t he bandit task better than three alternative models which differed in the expression of the neuromodulation component. Overall, these results validate the above mentioned hypotheses, confirming that the cholinergic influence on dopamine mediates uncertainty-seeking behaviors. Moreover, the model makes the following predictions which can be further tested experimentally: i) the correlation of dopaminergic activity with reward uncertainty as reported by Fiorillo et al. [2003] should not be observed in the absence of the cholinergic influence o n DA n eurons; ii) the adaptive advantage brought by the implemented uncertainty-seeking mechanism is most useful when sources of reward are not highly uncertain.

How animals generate variable decisions and manage the exploitation–exploration dilemma (i.e. choosing between predictably rewarding actions and other uncertain and suboptimal options) is still poorly understood. It has been suggested that humans rely on two types of exploratory behaviors [Wilson et al., 2014]: *directed exploration* in which uncertain actions are purposely chosen for the sake of information-gathering; and *random exploration* where actions are selected regardless of their predicted outcome. Our model formally describes how these two exploratory processes can be implemented: the former via the uncertainty bonus driven by the cholinergic influence on dopamine and the latter through a global decrease of dopaminergic modulation of decisions which results in lower selectivity and higher sensitivity to noise. This model could be thus be tested against other experimental data to further assess the validity of our hypotheses.

Moreover, some models suggested that the striatal cholinergic interneurons modulate the level of noise during action selection in the basal ganglia [Stocco, 2012]. This would imply a key role of acetylcholine, not only in directed exploration as we showed in this paper, but also in random exploration. We believe that new experimental studies are required which specifically investigate this possible dual implication of acetylcholine in exploratory processes. For instance, using tasks that leverage both random and directed exploration, lentiviral expression could selectively target cholinergic receptors in the striatum and in VTA to evaluate their respective involvement in these behaviors as well as possible interdependences. Furthermore, it is still unclear whether the cholinergic receptors in VTA dopamine neurons are required for learning the uncertainty bonus or solely for operating the bonus during action selection. These two alternatives could be differentiated experimentally via genetic-chemical manipulations rendering the cholinergic receptors light-controllable [Durand-de Cuttoli et al., 2018]. If the receptors are switched off during the initial sessions in which animals learn the statistics of reward delivery, and then switched on again, we should be able to find out whether the uncertainty seeking behavior appears rapidly or requires additional learning.

This work is a step toward a more comprehensive understanding of the implication of the dopaminergic and cholinergic systems in decision-making. It highlights their role in motivation and the execution of decisions. More effort is yet needed to further disentangle these neural mechanisms. For instance, more realistic neuron models could offer a complementary insight into the learning process [Deperrois and Gutkin, 2018]. It has also been suggested that to be able to account for both learning- and motivation-related processes, it is important to distinguish dopamine cell firing from local dopamine release on dopamine terminals [Berke, 2018]. Thus, a more detailed model of the decision-making network might be necessary to fully capture the role and functioning of the neuromodulators in these processes. By showing how ACh might drive uncertainty seeking behavior through its influence on DA, the present model is a first step in that direction.

## 6 Acknowledgements

The authors would like to thank Jérémie Naudé and the Neuroscience Paris Seine laboratory (Sorbonne Université, INSERM, CNRS) for sharing the data from their study [Naudé et al., 2016]. They are also grateful to Jérémie Naudé, Philippe Faure, Olivier Sigaud and Andrea Soltoggio for fruitful discussions and comments. MB is thankful to the ETIS laboratory for supporting his visit to UCI. At Sorbonne Université, MB is supported by the Labex SMART (ANR-11-LABX-65) which is funded by French state funds and managed by the ANR within the Investissements d’Avenir programme under reference ANR-11-IDEX-0004-02. JLK was supported in part by the United States Air Force and under Contract No. FA8750-18-C-0103. Any opinions, findings and conclusions or recommendations expressed in this material are those of the author(s) and do not necessarily reflect the views of the United States Air Force and DARPA.

## 7 Authors’ contributions

MB and JLK designed the study. MB implemented the model, analyzed the data and wrote the initial draft. MB and JLK reviewed and edited the manuscript.

